# An Algebraic Approach to Representing Mitochondrial Dynamics

**DOI:** 10.1101/2025.03.17.643721

**Authors:** Raphael Mostov, Greyson Lewis, Gabriel Sturm, Wallace F. Marshall

**Affiliations:** Dept. Biochemistry and Biophysics, University of California, San Francisco

**Keywords:** cell representation, algebraic graph theory, morpholomics

## Abstract

This paper addresses the increasing need for comprehensive mathematical descriptions of cell organization by examining the algebraic structure of mitochondrial network dynamics. Mitochondria are cellular structures involved in metabolism that take the form of a network of membrane-based tubes that undergo continuous re-arrangement by a set of morphological processes, including fission and fusion, carried out by protein-based machinery. Because of their network structure, mitochondria can be represented as graphs, and the morphological operations that take place in the cell, referred to as mitochondrial dynamics, can be represented by changes to the graphs. Prior studies have classified mitochondrial graphs based on graph-theoretic features, but an alternative approach is to focus not on the graphs themselves but on the set of morphological operations inducing mitochondrial dynamics, since this may provide a simpler representation. Moreover, the operations are what determine the graphs that will be generated in a biological system. Here we show that mitochondrial dynamics constitute a groupoid that includes the automorphism group of each mitochondria graph. For multi-component mitochondria we define a graph structure that encapsulates the structure of mitochondrial dynamics. Using one of the morphological operations we can define an equivalence relation among mitochondrial graphs that allows us to replace the complex structure of the full groupoid with a vastly simpler groupoid representation based on equivalence classes. Using these formalisms we define a distance metric for similarity between mitochondrial structures based on an edit distance. In the course of defining these structures we provide a mathematical motivation for new experimental questions regarding mitochondrial fusion, the impacts of cell division on mitochondrial morphology, and the presence of a single giant component in some cell types. This work points to a general strategy for formulating a cell structure state-space, based not on the shapes of cellular structures, but on relations between the dynamic operations that produce them.

## 1 Introduction

### 1.1 The problem of cell representation

Cell biology is drowning in data. With the advancement of high-throughput imaging and super-resolution methods, we are reaching a point where the quantity of information generated becomes difficult to comprehend. There is growing interest in developing mathematical approaches for representing cell structure in simpler terms that can be more tractable [1, 2, 3]. A formal way to represent cell structure would enable using the representation to classify cell types or cell states; to quantify similarity and differences between cell organization under different conditions; and to provide a way to describe the complex phenotypic effects of mutations.

Formal cell representations also provide a basis for mathematical models that can help us to understand, predict, and engineer cell behavior. The type of mathematical model used for analyzing cell behavior depends on the underlying representation used for cell state. The vast majority of mathematical models in biology have relied on representing biological processes using continuous variables and systems of differential equations [4]. Many biological questions remain difficult to model in terms of differential equations, suggesting a need for alternative mathematical modeling frameworks better suited to such questions. One alternative type of mathematical approach is algebra, in which a biological system is represented in terms of groups or other algebraic structures [5, 6, 7]. Here we develop an approach to use abstract algebra to represent the dynamics of a key cellular structure - the mitochondrion.

### 1.2 The Mitochondrion - an organelle with a network morphology

Mitochondria, often referred to as the “powerhouse of the cell,” are often depicted as bean-shaped objects in textbooks. But in reality, mitochondria in most cells exist as networks of tubules spread throughout the cell interior [8]. Mitochondrial network morphology is conspicuously different when cells are grown under different conditions [9], and also changes during the cell cycle [10], cell differentiation [11], and in various disease states [12]. The connectivity of mitochondria in a cell determines the ability of metabolic products to move within them [8, 13], as well as the ability of mitochondrial DNA to redistribute [14], and to be inherited [15, 16]. For all of these reasons, mitochondrial network morphology is biologically important, and the molecular mechanisms that produce these networks have been the topic of intense study.

What determines the network morphology of mitochondria [17]? Like any physical network, we can describe mitochondrial morphology using the mathematical definition of a graph with vertices and edges [18, 19]. For mitochondria, these graphs consist entirely of vertices of degree 1 or 3 [17]. Vertices of degree 4 (four-way junctions) sometimes occur transiently but rapidly resolve into pairs of degree 3 vertices. In some organisms, the network is constrained to lie on the surface of the cell, such that the graph is planar.

#### Definition 1.1.

A *mitochondria graph* is a graph composed exclusively of degree-one and degree-three vertices.

Several software packages, including MitoGraph [18] and Nellie [20], can be used to convert 3D images of mitochondria into graph representations (See Figure 1). This ability to render mitochondria as graphs makes it easy, in principle, to directly apply mathematical ideas from graph theory to actual biological data.

**Figure 1:**
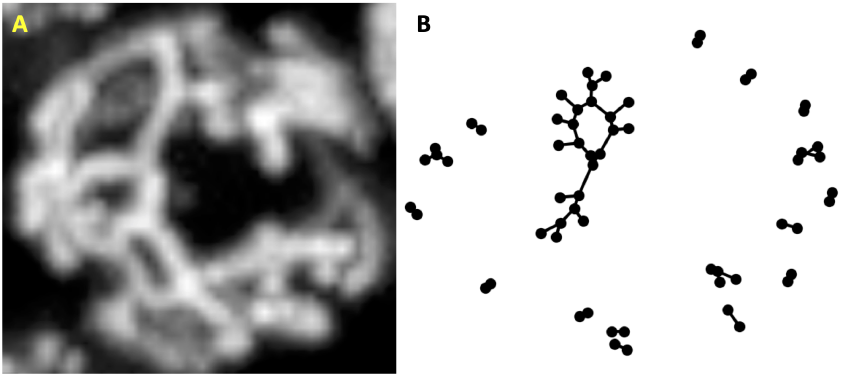
Graph representation of a mitochondria network. (A) max-intensity projection of 3D image of budding yeast cell expressing mito-dsRed, grown in glycerol and imaged by spinning disk confocal microscopy. (B) Mitochondrial graph structure of yeast cell in glycerol computed using MitoGraph 2 software [18].

### 1.3 Mitochondrial dynamics

One interesting feature of mitochondrial morphology is that the networks are dynamic, constantly changing their connectivity [21]. This occurs primarily through fission, the splitting of network branches, and fusion, in which two network branches join together [22]. There are two types of fusion: tip-to-tip, in which the ends of two different network branches join, and tip-to-side, where the end of one branch joins to the side of another. Additionally, there are several other morphological transformations; outgrowth: where a new branch grows out from the side of an existing branch; resorption: the inverse of outgrowth; mitophagy: the digestion and removal of small mitochondria; and the vertex flip, in which two vertices slide past each other due to branch migration, as diagrammed in Figure 2. As shown in Figure 2, we can describe these morphological transformations using the graph representation of mitochondrial networks [17]. Experimental studies show that altering the rates of these processes gives rise to mitochondrial networks with different graph structures [18].

**Figure 2:**
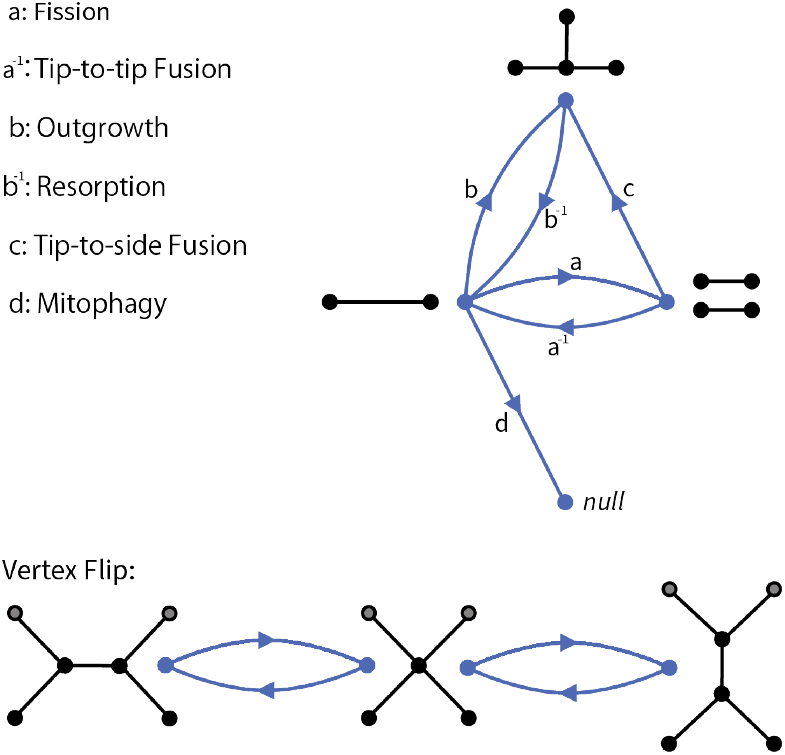
Morphological operations on mitochondria graphs.

By acquiring 3D images of mitochondria in live cells over time, and then converting the mitochondrial network to a graph at each time-point, it is possible to represent mitochondrial dynamics as a series of the above-listed morphological operations. Figure 3 shows one example from an actual cell in which a branch resorbtion event is followed by fusion of the large component with a smaller component, which is then followed by fission. Technology to acquire 3D images of mitochondrial networks in living cells at high spatial and temporal resolution, and to convert these images into graph representations, is by now quite well developed. But how can we gain insights from such rapidly expanding datasets?

**Figure 3:**
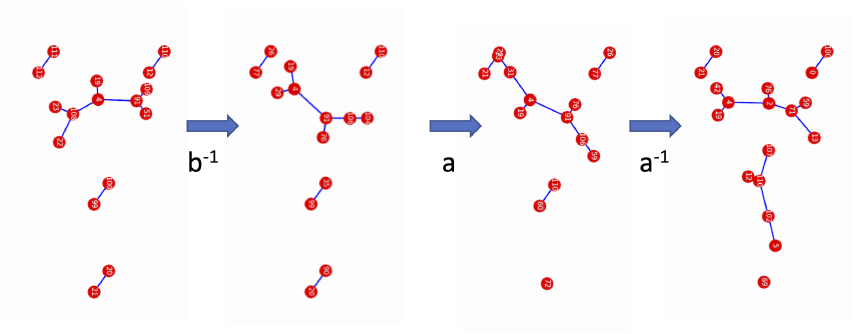
Graph dynamics in mitochondria in living cells. Images show four sequential 3D images of the mitochondrial network in a budding yeast cell. Strain: CGY49.77 wildtype with preCox4-mNeon-green & TIM20-mCherry, grown to stationary phase in synthetic complete medium supplemented with 2% glucose and 2% methyl-alpha-D-mannopyranoside, imaged by single objective light sheet microscopy at a rate of 1 3D volume per second, and graphs extracted using the Nellie software package [20].

Consideration of mitochondria as dynamically changing graphs raises a number of questions that are inherently mathematical in nature. What is the range of possible mitochondrial graphs, and what statistical distribution of graphs is expected as a function of the rates of the dynamic processes? How do the different processes contribute to the space of possible graphs? Is there any sort of control system in the cell that actively adjusts the choice of operations to tune the network structure, or are the various morphological operations acting independently and at random? Can we use the set of morphological operations to define a measure of similarity between mitochondrial graphs based on the number of operations needed to convert one to the other?

To answer these questions, it is necessary to understand the structure of the space of mitochondrial graphs and their transitions. However, the space of mitochondrial graphs is complicated, difficult to enuemerate due to the problem of graph isomorphism, and, from a theoretical standpoint, infinite. To date, the main approach to characterizing the range of mitochondrial graphs has been to apply graph theoretic descriptors like diameter or cyclomatic number to define statistical distributions of observed mitochondrial graphs, but the question remains, what processes determine the observed distributions? Here we propose to apply a different set of tools, drawn from abstract algebra, to think about mitochondrial network formation in a different way. Algebraic graph theory has seen only very limited application in biological problems, and has not to our knowledge been used to understand mitochondrial architecture. We believe that this work will furnish new mathematical problems while, at the same time, providing a new way to get at a fundamental biological question.

## 2 Results

### 2.1 State Space of Mitochondrial Graphs is Irreducible

In order to represent the space of mitochondrial graphs in terms of the set of morphological operations, it is necessary that the space be irreducible with respect to this set of transformations. We therefore first ask whether it is possible to find a sequence of the morphological transitions listed in Figure 2 that would allow us to interconvert between any two mitochondria graphs. The answer is yes. Notice, in Figure 4, that for any mitochondria graph, we can perform a series of fissions to transform it into a collection of Y-shaped tubules. Then, we can resorb one branch on each Y-tubule into an I-shaped graph, a mitochondria graph with only two degree-one vertices (i.e. *K*_2_), and tip-to-tip fuse the remaining connected components into a single mitochondrion. Since all of these transitions have inverses, fission being the inverse of tip-to-tip fusion and outgrowth the inverse of resorption, we can reverse this process to reach any other possible mitochondria graph. We recognize, of course, that in actual cells this sequence of events is highly unlikely to occur, but it is not impossible, and therefore we are justified in invoking irreducibility in derivations when necessary.

**Figure 4:**
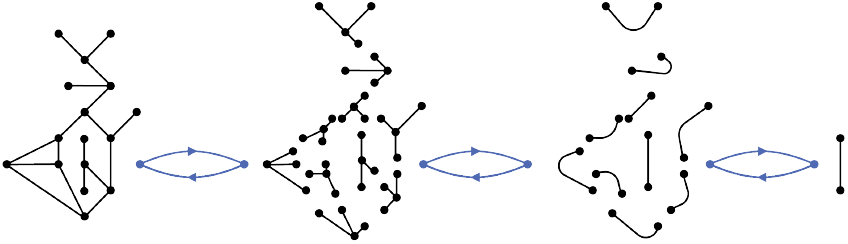
Visual explanation of how it is possible to convert from any mitochondria graph to a single mitochondrion and vice versa.

This simple result has immediate implications. Firstly, this implies that (a finite version of) the state space could, in theory, be modeled as a Markov chain with a stationary distribution, enabling predictions to be made using tools from statistical mechanics. Secondly, mitochondria have been observed to adjust their morphology similarly to the aformentioned thought experiment. The mitochondrial matrix has been observed to temporarily contract into nodules in a way that visually resembles a network fissioning into many smaller connected components – although the network remained connected by thin, almost invisible, tubules [23]. Surprisingly, Lee & Yoon [24] found that fission deficiency greatly amplified this phenomenon. This compaction into a chain of nodules is at least visually reminiscent of the proposed reduction to I-tubules. Third, and most relevant to our work, this interconvertability is a property of groups and several other types of algebraic structures, suggesting that there perhaps are hidden symmetries in mitochondrial dynamics. that could provide a new way to view relations between mitochondrial networks based not only on their traditional graph-theoretic properties but on relations between the transformations that produce them. The idea is to define an algebraic structure that describes how different morphological operations compose together, and then view the effect of these operations as actions on the set of mitochondrial graphs. In this viewpoint, the interconvertability property would correspond to a transitive action.

Mitochondrial dynamics offer a promising avenue to apply abstract algebra to raise questions of interest for both mathematics and cell biology. The key properties that one uses to define an algebraic structure have biological interpretations that could lead to meaningful insights. For instance, the interconvertability property of groups arises from the requirement that each element must have an inverse. Mitochondria are interconvertable, however, tip-to-side fusion does not have a direct inverse. Rather, as seen in Figure 2, tip-to-side fusion has a functional inverse that (in most cases) is obtained by first performing resorption followed by fission. In group theory terms, this functional inverse is a relation [25]. As implied in our interconvertability result, it is possible to interconvert between any two mitochondria graphs without tip-to-side fusion – it can be replaced by composing tip-to-tip fusion and outgrowth.

Knowing this, some questions immediately come to mind. Why does tip-to-side fusion exist if it is seemingly not necessary? If a mitochondrial network could not undergo tip-to-side fusion, what differences would we observe in its morphology? In their investigation of the mechanisms behind mitochondrial fusion, Gatti and colleagues [26] observed that about 75% of fusion events were tip-to-side. Furthermore, they found evidence suggesting that tip-to-side and tip-to-tip fusion are mechanistically different: actin was present in about 88% of tip-to-side fusion events but only in 50% of tip-to-tip fusion events. This example characterizing tip-to-side fusion demonstrates how the exercise of defining an algebraic structure on the mitochondrial dynamics state space encourages us to consider mitochondrial biology from new perspectives.

### 2.2 Mitochondrial Network Dynamics forms a Groupoid

The requirement that a group’s multiplication be a total binary function, defined for all pairs of elements in the group, is not consistent with our representation of mitochondrial network dynamics as graph operations, with composition of morphological operations as multiplication, because it is not necessarily the case that operations can be composed. For example, it is possible for a branch resorption to occur on the sole pendant edge of a graph, such that tip-to-tip fusion is no longer a valid operation on the resulting graph. Thus, it is not the case that any two operations can be combined, hence the multiplication (composition) is only a partial binary function. A more appropriate framework would be one that satisfies all of the group properties except for the requirement of a well-defined group multiplication – namely a groupoid, which is a set with an operation defined as a partial binary function to which we can’t necessarily apply the operation to any arbitrary pair of elements. For the sake of rigor, we also provide a formal definition [27].

#### Definition 2.1.

A *groupoid* consists of a set of elements *G* together with a set of objects Ob(*G*) as well as source and target maps *s, t* : *G* → Ob(*G*) such that the following properties hold for all *x, y, z* ∈ *G*:

- *Partial Multiplication*: If *t*(*x*) = *s*(*y*), then the product *xy* ∈ *G* is defined.
- *Associativity* : If the product *xyz* ∈ *G* is defined then *xyz* = (*xy*)*z* = *x*(*yz*).
- *Inverses*: There exists *x*^−1^ ∈ *G* such that the products *xx*^−1^ and *x*^−1^*x* are defined, and…
- *Identity* : If *xy* ∈ *G* is defined, then *xyy*^−1^ = *x* and *x*^−1^*xy* = *y*.

One of the most familiar examples of a groupoid is the 15-puzzle consisting of tiles that can slide around to rearrange their order (Figure 5(A)). For any configuration of the 15-puzzle, there are between two and four types of transformations we can apply to the puzzle. As seen in Figure 5(B), if the empty square is not on the edge of the puzzle or in a corner, we can move it up, down, left, or right. Here, Ob(*G*) is the set of all possible configurations of the puzzle, of which there are 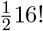 [28], and *G* is the set of all possible transformations (movements of tiles). We consider two transformations to be the same groupoid element if they have the same starting and ending configuration – the same source and target. For each groupoid element, an inverse element exists, defined by the opposite transformation. Furthermore, there is an identity element, which corresponds to the ‘do nothing’ transformation.

**Figure 5:**
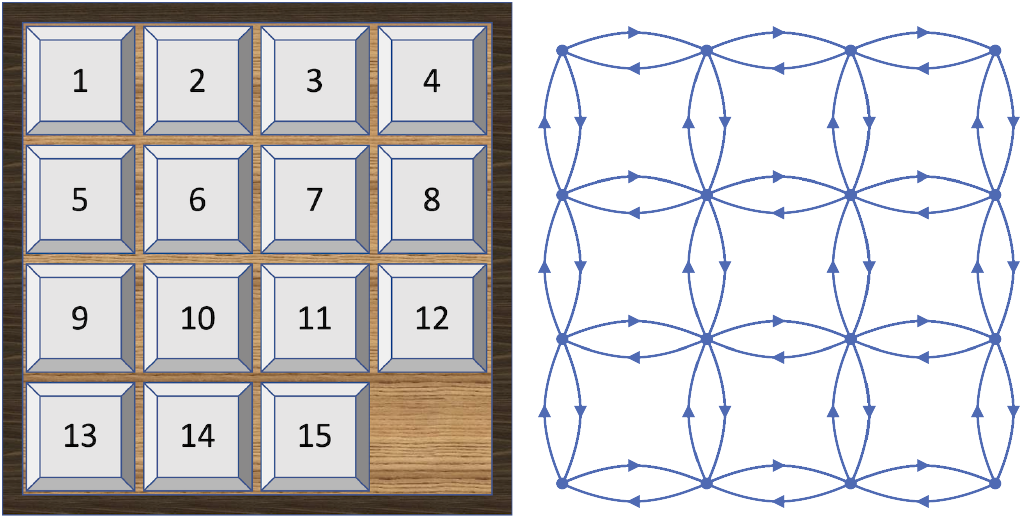
(Left) Solved position of the 15-puzzle. (Right) Graph depicting the possible locations and movements of the empty tile in the 15-puzzle.

The groupoid formalism has a clear interpretation as a discrete state space. We can view each element of Ob(*G*) as a state and each groupoid element as a transition from a source state to a target state. From this perspective, we can define a sense in which mitochondrial network dynamics form a groupoid.

#### Definition 2.2.

The *mitochondria groupoid MG* is the groupoid in which Ob(*MG*) is the set of mitochondria graphs, and the elements of *MG* are the morphological transitions of mitochondrial dynamics.

The elements of *MG* are not just {*a, a*^−1^, *b, b*^−1^, *c, d, e*}. Rather, each element is specified by its transformation, as well as source and target mitochondria graphs. For instance, we can have an element *a* : *M* → *M′* that fissions a specific edge in *M* to create *M′*, but we can also have another element *a* : *M* → *M′′* that creates a different graph by performing fission on a different edge.

With this caveat in mind, one can further check that all the state space transitions (morphological operations), except *c*, have direct inverses and are thus valid groupoid elements. The inverse of *c* is obtained by performing *ab*^−1^ on the appropriate edges. Many important mathematical results that make use of groups, such as Van Kampen’s Theorem or the Burnside Lemma, have generalizations that extend to groupoids [27, 29]. While the groupoid *MG* that we have outlined encompasses the full family of mitochondria graphs, its current description is likely too abstract to apply any relevant results. As such, we will spend the rest of this section trying to make the descriptor of mitochondria graphs more tractable while still having a groupoid that is comprehensible.

### 2.3 A Simplified Representation of Mitochondria Graphs Based on an Equivalence Relation

The big challenge with cell representation in general, is the large size of the possible space of cell structures. This is exemplified by the problem of trying to enumerate the space of possible mitochondrial graphs. As the graph becomes large, the number of possible graphs, as well as the number of distinct graphs that can be produced from a given graph by a set of transformations, grows combinatorially. For example, it is often possible to apply a particular operation such as fission to many possible edges in a graph to produce different graphs as an outcome. How can we simplify the problem? One approach is to find some criterion by which collections of mitochondrial graphs can be viewed as equivalent.

Among the morphological operations defined above, vertex flip can uniquely be used to define an equivalence relation, which we define here and justify below.

#### Definition 2.3.

We will say that mitochondria graphs *M* and *M′* are vertex flip equivalent if there exists a sequence of vertex flips taking *M* to *M′* and vice versa.

In contrast to the other operations, vertex flip does not alter the number of degree 1 or 3 vertices, hence for two graphs to be vertex flip interconvertable, the number of degree 1 and 3 vertices must be the same for both graphs. We can keep track of the number of these quantities with the vector *M* = [*p, n*], in which *p* and *n* are the number of degree 1 and degree 3 vertices respectively. Before exploring how morphological operations work in this representation, we note that there are constraints on this representation such that only certain combinations of *p* and *n* are possible. First, the number of vertices *p* or *n* cannot be less than zero. Second, given the value of n, the value of p must obey defined bounds:

#### Proposition 2.1.

*Let* [*p, n*] *be the vector representing a single connected component of a mitochondria graph, then the maximum value of p is n* + 2 *and the minimum is* 0 *if n is even and* 1 *if n is odd*.

*Proof*.

1. (Upper Bound) Consider the induced subgraph *Y* ⊆ *M*, where the set of vertices *N*_*Y*_ of *Y* are all the degree-three vertices in *M*, and *n* = |*N*_*Y*_ |. Note that the degree of a vertex in *Y* is not always three, but can be one, two, or three depending on the other vertices in *Y* it’s connected to. Since there are *p* degree-one vertices in *M* which collectively are adjacent to the set of *n* vertices in *Y*, it follows by definition of *M* and *Y* that 3*n* = *p* + ∑ _*n*_ deg(*y*_*n*_). By the Handshaking Lemma, we have *p* = 3*n* − 2|*E*_*Y*_ |. Hence to maximize *p*, we must minimize the number of edges in *Y*. From Proposition 1.3.13 in [30], the minimum is *n* − 1. Thus, the maximum of *p* is *p* = *n* + 2.
2. (Lower Bound) Consider the cases in which *n* = 1 and *n* = 2. For the *n* = 1 case, it follows from Proposition 1.3.5 in [30], that we must have an even number of odd-degree vertices, and hence we must have another odd-degree vertex that is not a degree-three vertex. Hence the minimum amount of degree-one vertices for the *n* = 1 case is *p* = 1 (this graph would have a single self-loop and be shaped like the letter “P”).

For the second base case, *n* = 2, we can reach it by performing outgrowth on the graph given by *n* = 1, and as a consequence, we must produce a new degree-one vertex. We can then fuse the two degree-one vertices together into a single edge resulting in a graph with *p* = 0 degree-one vertices for the *n* + 1 case. We can inductively continue this process of performing outgrowth, and then performing tip-to-tip fusion as soon as two degree-one vertices are available..

Second, we note:

#### Proposition 2.2.

*For any connected component of a mitochondria graph* [*p, n*], *p can be any element of the set p* ∈ {0, 2, 4, …, *n, n* + 2} *if n is even and p* ∈ {1, 3, 5, …, *n, n* + 2} *if n is odd*.

*Proof*. Consider the base case with *p* = 0 if *n* is even and *p* = 1 if *n* is odd. We can perform fission on an edge while maintaining a single connected component to get [*p* + 2, *n*]. We can repeat this process until we get [*n* + 2, *n*], at which point we can’t maintain a single connected component.

Now we can return to the concept of vertex-flip equivalence. The following proposition shows that all single-component graphs with the same number of degree-three and degree-one vertices are vertex-flip equivalent.

#### Proposition 2.3.

*Let M and M′ both be single-component mitochondria graphs with the vector representation* [*p, n*], *then M and M′ are vertex flip equivalent*.

*Proof*. We would first like to note that it is cumbersome to define a vertex flip without using a picture. To make this proof easier to understand and follow, we will rely heavily on pictures without using any other definition of a vertex flip. Our proof relies on induction, and we have two mitochondria graphs which serve as base cases:

- The vector [0, 0], is the null mitochondria graph which corresponds to a toroidal shaped mitochondrion. It is vacuously vertex flip equivalent.
- The vector [2, 0] represents only one possible mitochondria graph, an I-tubule. It is vacuously vertex flip equivalent.

Assume inductively that all graphs with a single connected component described by [*p, n*] are vertex flip equivalent. Notice in Figure 6 that we can move one pendant edge next to any other edge with just vertex flips by “sliding” the pendant edge along its “neighboring edge”. Since we get [*p* + 1, *n* + 1] by outgrowing a pendant edge from [*p, n*], and can get back to a [*p, n*] graph by resorbing a pendant edge, then all graphs [*p* + 1, *n* + 1] are vertex flip equivalent. Moreover, we can obtain [*p, n* + 2] from [*p, n*] by inserting a new edge such that it connects two edges (by performing two outgrowth steps followed by a tip-to-tip fusion step).

**Figure 6:**
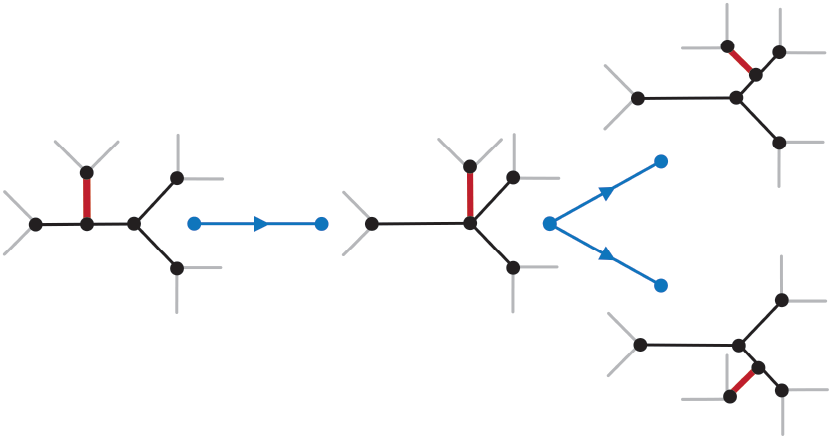
Image showing how it is possible to move the red colored edge next to any other edge using only vertex flips by “sliding it along” its “neighboring” edge.

As shown in Figures 6 and 7, we can move this edge next to any other edge using only vertex flips by “sliding” the edge along its neighboring edges. Hence all graphs [*p, n* + 2] are vertex flip equivalent. It follows from induction and proposition 2.2 that the set of graphs corresponding to any [*p, n*] are vertex flip equivalent.

**Figure 7:**
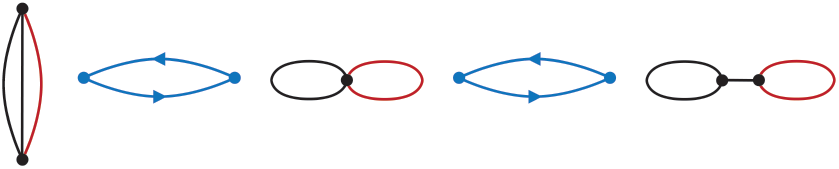
Image showing how both the mitochondria graphs represented by [0, 2] are vertex flip equivalent.

Finally, we note several properties of vertex flip equivalence.

1. First, it is straightforward to show that any graph is vertex flip equivalent to itself.
2. Second, if graph *M′* can be reached from graph *M* by a sequence of vertex flips, then the converse is also true.
3. Third, if graphs *A* and *B* are vertex flip equivalent, and graphs *B* and *C* are vertex flip equivalent, then so are graphs *A* and *C*.

Because the relation of vertex flip equivalence is reflexive, symmetric, and transitive, it is an equivalence relation, hence the set of all graphs that are vertex-flip equivalent to any given graph constitute an equivalence class. Each distinct vector [*p, n*] defines one such equivalence class, which taken together partition the space of possible single-component mitochondrial graphs. We note that none of the other operations of mitochondrial dynamics (fission, fusion etc) satisfy the symmetry property needed to define an equivalence relation.

We can take this representation further. If a mitochondria graph has multiple connected components, then we can represent each connected component with a vector, and the resulting multiset of vectors will be it’s own equivalence class.

#### Definition 2.4.

A *multiset vector representation* of a general mitochondria graph is a multiset of vectors in which each vector counts the number of degree-one and degree-three vertices in a connected component. For instance,

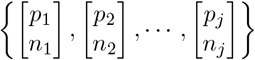

represents a mitochondria graph with *j* connected components.

### 2.4 Mitochondria Groupoid Can be Simplified With Vertex-Flip Equivalence Classes

For the multiset vector representation, we must define the transformations that change the number of connected components differently from those that don’t. For instance, if we have the graph representation [3, 3] and we want to perform fission on it, there would be two ways of doing so. We could fission a non-pendant edge to get the graph represented by [5, 3], or we can fission a pendant edge to get {[3, 3], [2, 0]}.

More generally, fission on any edge that is a cut-edge would result in addition of a new vector to the multiset. We must therefore define a ‘connected fission’ *a*_1_ which maintains the number of connected components, and a ‘disconnected fission’ *a*_2_ which increases the number of connected components. Similarly, we must define connected and disconnected fusions (see Figure 8).

**Figure 8:**
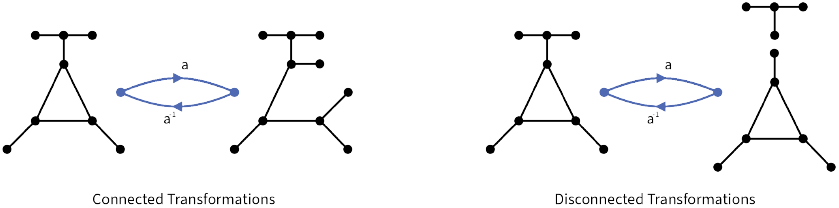
Comparison of connected fission/tip-to-tip fusion with disconnected fission/tip-to-tip fusion.

The set of morphological operations defined on the space of vector multisets can then be represented as follows:

- *Connected Fission*: *a*_1_([*p, n*]) = [*p* + 2], *n*, where *p < n* + 2.
- *Connected Tip-to-Tip Fusion*: 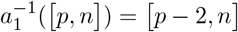, where *p* ≥ 2.
- *Disconnected Fission*: *a*_2_([*p, n*]) = {[*p*_1_, *n*_1_], [*p*_2_, *n*_2_]}, where *n*_1_ + *n*_2_ = *n, p*_1_ + *p*_2_ = *p* + 2, and *p*_1_, *p*_2_ ≥ 1.
- *Disconnected Tip-to-Tip Fusion*: 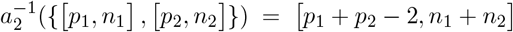, where *p*_1_, *p*_2_ ≥ 1.
- *Outgrowth*: *b*([*p, n*]) = [*p* + 1, *n* + 1]
- *Resorption*: *b*^−1^([*p, n*]) = [*p* − 1, *n* − 1], where *n, p* ≥ 1.
- *Connected Tip-to-Side Fusion*: *c*_1_([*p, n*]) = [*p* − 1, *n* + 1], where *p* ≥ 1.
- *Disconnected Tip-to-Side Fusion*: *c*_2_({[*p*_1_, *n*_1_], [*p*_2_, *n*_2_]}) = [*p*_1_ + *p*_2_ − 1, *n*_1_ + *n*_2_ + 1], where *p*_1_ ≥ 1 and/or *p*_2_ ≥ 1.
- *Mitophagy*: *d*([2, 0]) = ∅, only possible on [2, 0].

These operations are defined on individual vectors or pairs of vectors in a multiset, which are then replaced with the specified new vectors while all other vectors in the multiset are kept unchanged. No known mitochondrial morphological operations act on more than two components.

The restrictions on these transformations give us:

#### Definition 2.5.

The *small mitochondria groupoid mG* is the groupoid in which Ob(*mG*) is the set of multiset vector representations of mitochondria graphs as defined above, and the elements of *mG* are the sets of transitions between Ob(*mG*). Furthermore, there is a different version of *a*_2_ defined for each possible way there is to split up a single connected component into two connected components.

As was the case for the mitochondria groupoid *MG*, we also note here that the elements of *mG* are not 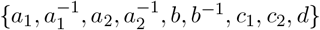, but rather each such transformation may constitute a different groupoid element depending on which vector(s) it is performed on.

We see that *mG* constitutes a mathematical model for mitochondrial dynamics in a simplified state space based on equivalence classes of mitochondrial structures. Notice that *mG* is obtained from *MG* by simply applying the vertex flip equivalence relation.

### 2.5 Mitochondria Groupoid Maps Onto a Discrete Vector Space

The mitochondria groupoid *mG* simplifies the representation of mitochondrial graphs by representing them as multisets of vectors corresponding to equivalence classes of graphs. This representation, while simpler than the groupoid defined directly on mitochondrial graphs *MG*, is still extremely complex. For each object in *mG*, there are a large number of groupoid elements corresponding to all the possible morphological operations operating on each of the components contained within a multiset of vectors, and which can either change one vector to another or else alter the number of components. We will next consider two possible ways of further simplifying the complex structure of *mG* to yield a more tractable representation at the expense of some detail or generality.

The first approach we consider for simplifying the mitochondria groupoid *mG* is by choosing to represent each mitochondria graph, including graphs with multiple components, as a single vector, rather than a multiset of vectors,.

#### Definition 2.6.

A *vector representation* of a mitochondria graph *M* is a vector [*p, n*] where *p* and *n* are the total number of degree-one and degree-three vertices respectively in the mitochondria graph.

By ignoring the difference components of each graph, it is no longer the case that all graphs described by a given vector are vertex-flip equivalent, because they can have different numbers of components. However, using a single vector admits a simplification of the mitochondria groupoid into a vector space. We can now define each morphological transformation simply as the addition of constant vectors:

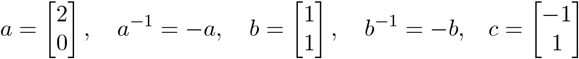

Composition of transformations is thus achieved by vector addition. With this representation we can represent sequential changes in the graph structure of real mitochondria observed in live-cell images in terms of sequential vector addition operations. Using these operations it is possible to describe a mitochondria graph as the sum of transitions through which it can be reached from some reference point, say the graph represented by [2, 0] (which we will denote as *I*). For instance, we can represent a mitochondria graph as the vector [2, 4] or as *I* + 2*b* + 2*c*. This discrete vector space defines another groupoid:

#### Definition 2.7.

The *vector groupoid vG* is the groupoid in which the objects Ob(*vG*) are the single-vector representations of mitochondria graphs and the elements are generated by the vector versions of the transitions *a, b, c, a*^−1^, *b*^−1^ subject to the constraint *c* = *ba*^−1^.

A major advantage of *vG* compared to either *MG* or *mG* is that it can be easily depicted in a two dimensional diagram. This is not the case for the earlier representations whose structure is difficult to visualize.

The simple two-dimensional lattice structure of this simplified representation makes it straight-forward to visualize the distribution of mitochondrial graphs structures in actual data. As shown in 9 panel B, we computed the number of degree 1 and 3 vertices from mitochondrial graph data reported in [18] who used 3D imaging of budding yeast cells followed by application of MitoGraph software to generate graph representations of mitochondria in individual cells. Comparing the real data to the diagram of *vG* shows a broad distribution of vectors seen in real cells suggesting that the full state space is accessed. By using the simple representation of *vG* we gain the ability to easily map observational data onto our representation at the cost of losing the ability to invoke vertex flip equivalence.

Because it is defined with objects as vectors rather than graphs, this groupoid merely has the constraint that we cannot produce a graph with a negative number of vertices.

### 2.6 A Distance Metric for Mitochondria Graphs

Another application of the *vG* and *mG* models is that we can use them to define, within *vG* and *mG* respectively, a distance between any two mitochondria graphs in the state space, in terms of the path with the minimum number of transitions, which could be used to cluster mitochondrial with similar graph structures, estimate the time required to go from one structure to another, and so forth. An important caveat is that because tip-to-side fusion does not have a direct inverse, these distance metrics are asymmetric; that is, the distance from A to B is not necessarily the same as the distance from B to A.

#### Proposition 2.4.

*Let* [*p, n*], [*p*^*′*^, *n*^*′*^] ∈ *vG be vector representations of mitochondria graphs. The* ***asymmetric taxicab distance*** *is*

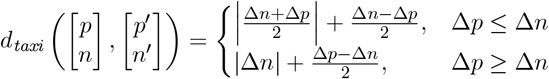

*where*

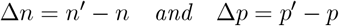

*Proof*. There exists *α, β* ∈ ℤ and *γ* ∈ ℕ such that

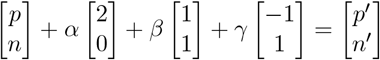

the taxicab distance is

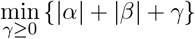

constrained by the above linear equation. We thus solve the minimization problem.

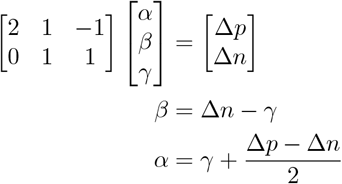

and so

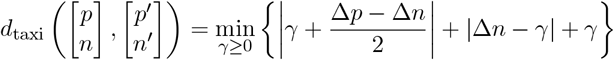

Since this is a piecewise linear convex function, the minimum must occur either at the boundary *γ* = 0, or, at a place where one of the absolute value terms is 0 – either when *γ* = Δ*n*, which is feasible when Δ*n* ≥ 0, or 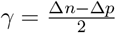, which is feasible when Δ*p* ≤ Δ*n*. After checking each of the possibilities, we arrive at the asymmetric taxicab distance defined in Proposition 2.4

Unfortunately, we are currently unable to find a similar distance for the representation *mG*. Actually finding the minimum is difficult because of the need to specify which connected components any given transformation will operate on. This can however lead to an algorithm that searches a constrained subset of all possible paths between two mitochondria graphs in order to determine the exact distance. Such an algorithm may have a time complexity that is small enough to be implemented on a computer. At the very least, we can obtain an upper and lower bound on the distance between any two mitochondria graphs that only requires us to know the multiset vector representation used in *mG*. We start with the lower bound.

#### Proposition 2.5.

*Let M and M′ be mitochondria graphs where n, n*^*′*^ *are the total number of degree-three vertices, p, p*^*′*^ *are the total number of degree-one vertices, and C, C′ are the number of connected components respectively. If C′* ≥ *C, then*

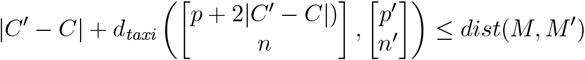

*If C′ < C, then*

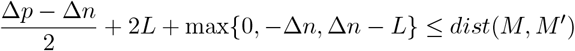

*where*

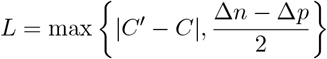

*Proof.*

- If *C′* ≥ *C*, then the only possible way to produce the necessary additional connected components is through |*C′* −*C*| additional fission steps. Thus, we first perform |*C′* −*C*| fission steps, and then compute the taxicab distance from that point to obtain the lower bound.
- If *C′ < C*, then the only way to get rid of the excess connected components is through |*C′* − *C*| additional tip-to-tip and/or tip-to-side fusion steps (in this instance, mitophagy can be counted the same as tip-to-tip fusion on an I-tubule). Thus, similar to our proof of Proposition 2.4, we minimize

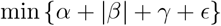

subject to the constraints

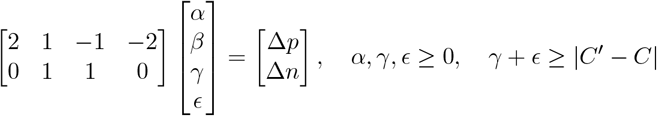

solving the linear equation constraint for *α* and *β* we get

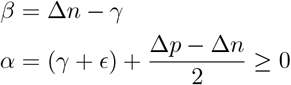

Hence,

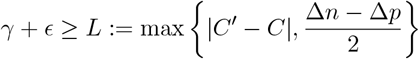

and our objective function becomes

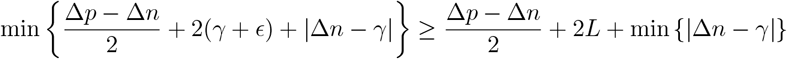

Let *s* = *γ* + *ϵ*, then, for fixed values of *s*, |Δ*n* − *γ*| is minimized by taking *γ* to be the integer in [0, *s*] closest to Δ*n*,

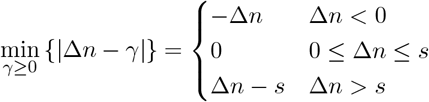

Minimizing *s* ∈ ℕ over *s* ≥ *L*, we arrive at the lower bound in Proposition 2.5.

Next, we use the information from the *mG* representation to derive on upper bound on the mitochondria graph edit distance.

#### Proposition 2.6.

*Let M and M′ be mitochondria graphs where n, n*^*′*^ *are the total number of degree-three vertices, p, p*^*′*^ *are the total number of degree-one vertices, and C, C′ are the number of connected components respectively. Then*

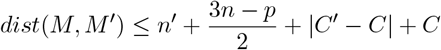

*Proof*. To obtain the upper bound, we find an upper bound on the distance from *M* to an intermediate graph, *H*^*′*^, which is composed of *C′* I-tubules, and then find the exact distance from *H*^*′*^ to *M′*. By the triangle inequality, *dist*(*M, M′*) ≤ *dist*(*M, H*^*′*^) + *dist*(*H*^*′*^, *M*).

1. Distance from *M* to *H*^*′*^: Let [*p*_*i*_, *n*_*i*_] be a connected component in *M*. Notice from Proposition 2.1 that

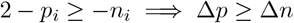

and so, since no matter where on any of the graphs represented by [*p*_*i*_, *n*_*i*_] we perform the transitions to get to the graph represented by [2, 0], we will end up at an I-tubule, then

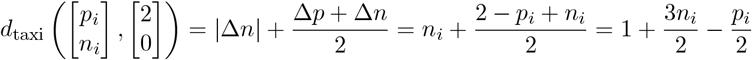

is the exact edit distance between [*p*_*i*_, *n*_*i*_] and [2, 0]. If *C′* ≥ *C*, then the only way to produce the additional connected components is through |*C′* − *C*| additional fission steps. If we first transform *M* into the graph with *C* I-tubules, then perform the extra fission steps, we will end up with *C* I-tubules. Similarly, if *C < C′* then we must perform at least |*C′* − *C*| additional fusion or mitophagy steps to end up with *C* connected components. Since performing tip-to-side fusion on two I-tubules makes one I-tubule, we can follow the same process as before to get

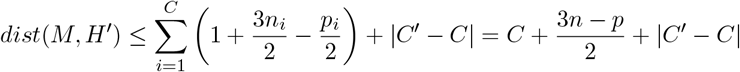
2. Distance from *H*^*′*^ to *M′*: Let [*p*_*j*_, *n*_*j*_] be a connected component in *M′*. By Proposition 2.1, we have

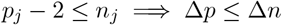

and so

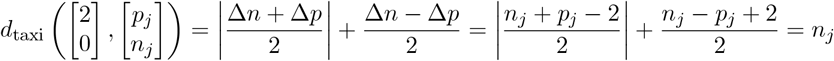

is the exact edit distance. Hence,

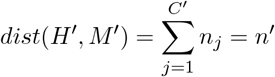

### 2.7 A Mitochondria Groupoid for Graphs with a Single Connected Component

An alternative way to simplify the complex structure of *mG* is to notice that much of the complexity arises from the fact that the objects are multisets of vectors, which requires us to keep track of operations that change the number of components. If we consider only objects from *mG* with a single component, and only elements that correspond to connected transformations (those that do not change the number of components), this amounts to introducing a constraint on *vG* from Proposition 2.1 that *p* ≤ *n* + 2. In this case, we retain the ability to easily map observed graphs onto vectors just by counting vertices of degree 1 and 3, but now the vectors once again represent equivalence classes of mitochondrial graphs based on vertex-flip equivalence (see 2.3), this time restricted to graphs with a single component. Thus, we have:

#### Definition 2.8.

A *single-component vector representation* is a representation [*p, n*] of a mitochondria graph *M* which has only one connected component, where *p* is the number of degree-one vertices and *n* is the number of degree-three vertices.

#### Definition 2.9.

The *single-component groupoid mmG* has objects Ob(*mmG*) that are single-component vector representations, and elements *mmG* that are identical to that of *vG*.

The Cayley Graph of *mmG* is described in Figure 10. The structure of *mmG* arises from the fact that within this representation, a single operation can only be applied in one way, and the action of any operation on vertex-flip equivalent graphs in the vector produces vertex flip equivalent outputs, as illustrated in Figure 11.

By using *mmG* to represent the structure of mitochondrial dynamics, we gain mathematical tractability as well as a straightforward way to map observed data onto the representation, but we do so at the cost of making an apparently extreme assumption that the mitochondrial network consists of a single connected component. This may at first seem to be a drastic assumption. In reality, budding yeast mitochondria often do in fact take the form of a single large connected component plus some number of I tubules [18]. A similar situation has also been seen in other cell types [31]. These are all still multi-component graphs. However, in such cases, fusion with an I tubule would either be by tip-to-tip fusion, in which case it would simply add onto the end of a pendant edge and thus have no effect on the graph structure of the giant component, or else it would fuse with the giant component by tip-side fusion in which case it would mimic branch outgrowth which is already one of the elements of *mmG*. A fission operation that produces a new I tubule would have to have involved a pendant edge, and leaves behind a pendant edge in the same position, so again it would not affect the graph structure of the giant component. So, the presence of a collection of I tubules along with a single giant component has no effect on the algebraic structure of the transformations operating on the single giant component. Thus, depending on cell type and growth condition, the single-component mitochondrial groupoid *mmG* may in fact be a biologically reasonable approximation of real mitochondrial dynamics.

## 3 Discussion

### 3.1 Mathematical questions raised by this study

In this study, we have only presented a method for building a representation of mitochondrial dynamics using algebraic methods. We hope that this work may suggest new areas for investigation by mathematicians. Here, we discuss several such potential directions:

#### 3.1.1 The Mitochondrial Groupoid MG Contains the Automorphism Group of Each Mitochondria Graph

The mitochondrial structure graph *MG* described here is evidently a complex structure. One approach to understanding this system is to look for structures that are contained within the larger structure.

Consider all the transformations that take a graph *G* in *MG* back onto itself. For any such transformation, its groupoid inverse is also part of the set. When only transformations from G to G are considered, the product becomes a binary function, unlike the general case of transformations between graphs. All other properties (associativity, inverse, identity) are unchanged, and hence the set of such transformations constitutes a group. One way to characterize the structure of *MG* will, thus, be to characterize these groups of return transformations for each mitochondria graph *G* of *MG*.

Such an approach has previously been used to investigate the algebraic structure of the 15-puzzle. In addition to being a groupoid, the puzzle can also be viewed as a group when we restrict ourselves to all configurations that have the empty square in a fixed position. If we fix the empty square in say, the bottom right corner, then the 15-puzzle becomes a permutation group since each possible tile configuration is just a permutation of the tiles from their solved position [32]. Every permutation can be written as a sequence of transpositions. We obtain the 15-puzzle group by allowing the empty square to move freely and imposing the restriction that we can only perform a transposition if it represents a legal sliding move. A notable result is that the 15-puzzle group is isomorphic to the group of all even permutations on 15 elements, where an even permutation is one that can be expressed as a sequence of an even number of transpositions [32, 33]. We expect that a similar approach may help clarify the structure of MG.

We therefore consider the set *Q*_*G*_ of strings of transformations (i.e. groupoid elements generated by the morphological operators) *q* that start and end at a particular mitochondria graph *G*. If we label each vertex in *G*, then some choices of *q* could permute the vertex labels. In the same way that the sliding move sequences in the 15-puzzle that keep the empty tile in the bottom right corner form a permutation group, so too could *Q*_*G*_. To properly define a group on *Q*_*G*_, we would have to determine how each morphological transformation affects the vertex labels of a mitochondria graph. Some transformations, like fission and outgrowth, add new vertices, while others, like tip-to-tip fusion and resorption, remove them – how would we add or remove the corresponding vertex labels? In fact it is possible to add and remove vertex labels in such a way that any choice of *q* would not add or remove vertex labels from *G*, but rather just permute the labels.

##### Proposition 3.1.

*Let G be a a mitochondria graph with N vertices, and consider the following labeling scheme:*

- *Label the vertices with natural numbers* 1 *through N by arbitrarily assigning the labels*.
- *Whenever we add a vertex to G, we label the new vertex N* + 1.
- *Whenever we remove a vertex numbered k from G, if k < N, then we relabel vertex N to k*.

*Furthermore, let Q*_*G*_ *be the set of strings of transformations that start and end at G. Then Q*_*G*_ *is the permutation group on N elements, and, contains the automorphism group of G*.

*Proof*. Let *G* be a mitochondria graph with *N* vertices. Taking note of our explanation of how mitochondrial dynamics are fully interconvertable back in section 2.1, we can first convert *G* into a graph *G*^*′*^ that consists of *N* I-tubules such that each I-tubule contains a vertex *k* ∈ {1, …, *N*} connected to a vertex *k* + *N*.

Next, consider the vertices labeled *i* and *j* in *G*^*′*^ where *i, j* ≤ *N*. If we want *i* to be a pendant vertex and a neighbor of *j* in the final graph, then we tip-to-tip fusion vertices *i* + *N* and *j* + *N* together. Else, if we want *i* to be a degree-three vertex and a neighbor of *j*, then we can tip-to-side fusion *i* to the edge connected to *j*. Iterating this process across all *N* of the I-tubules in such a way that we end back up at the graph *G*, we see it is possible to reach any relabeling of *G*. Thus, there are no “illegal configurations” in *Q*_*G*_, and *Q*_*G*_ is the permutation group on *N* elements, which must contain the automorphism group of *G* as a subgroup [34].

Can this type of approach be extended to the simplified groupoids *mG, vG* or *mmG*? Considering the vectors or multisets of vectors as objects, the collection of elements that return a vector M back to itself constitute a group as they would for any groupoid. These groups are different for different vectors because the constraints on valid vectors (see for example Figure 10) mean that some strings of transformations that can be applied to one vector M cannot be applied to another without producing an invalid vector. A related question is to ask if the automorphism groups of the graphs corresponding to a particular vector representation might tell us anything about the structure of the group of return transitions for the vector. In this context we note that it is easy to show that vertex flip equivalent graphs do not necessarily have the same automorphism group. A simple example is provided by the vector [4, 0] which includes the complete graph *K*_4_, for which the automorphism group is the symmetric group on four elements *S*_4_, the square graph with two multiedges, for which the automorphism group is the dihedral group on a square *D*_4_, as well as two other graphs, both containing self-loops, for which the automorpishm group is the integers modulo two, ℤ_2_ (see Figure 12).

#### 3.1.2 Planarity

In some cells, such as budding yeast, the mitochondrial network is constrained to lie on the surface of the cell, hence the mitochondrial graphs are plane graphs. How will the groupoid structure be changed by imposing the condition of planarity on the underlying graphs? For MG, planarity will entail removing any non-planar graphs from the set of objects. For the simplified groupoids based on the vector representation, can we be sure that one or more planar graphs exist for every [*p, n*] vector? To see that this is the case, note that the addition of pendant edges (increasing *p*) does not affect whether a graph is planar or not. It is thus sufficient to enumerate the planar 3-regular graphs through the process described in the proof of Proposition 2.1.

Will claims about vertex flip equivalence still hold if the underlying graphs are restricted to being planar? We believe this is an area for further exporation. For example, it is not yet clear if we can use vertex flips to migrate a pendant edge so that it is embedded in a different face.

### 3.2 Biological questions and applications

We believe that this work, while abstract in nature, may have direct applications in biology in three distinct ways. First, the process of making such simplified representations forces us to make our assumptions explicit, raising questions that we had not previously explored. One example of a question raised by the effort of simplifying algebraic structures is the question of how a mitochondrial network becomes partitioned during cell division. Second, the simplified framework helps us to ask about how how mathematical constraints may shape the observed biological structures. For example, what are the implications of tip-side fusion which produces an asymmetry in the Cayley graph; why is there often a single giant component in many cell types; and how might we test for morphological homeostasis. Finally, as with any case of cell representation, we believe that the simplified representation employed here can serve as the basis for a statistical methodology to define and compare cell states based on mitochondrial images by allowing us to assign a pair of integers, or a multiset of such pairs, to each graph structure.

#### 3.2.1 How do mitochondria graphs partition during cell division?

One benefit of building mathematical models in biology is that the mere act of formalizing a model forces the researcher to make their biological assumptions explicit, which can often reveal gaps in knowledge. One example in the case of mitochondria has to do with deciding on where to impose a finite constraint on the size of *MG* (or any of the simplified versions). While the structure grows forever as new vertices and edges are added to the mitochondrial graphs, at some point, the cell divides. Thus, the infinite size of *MG* means it could model an ensemble of mitochondria graphs across multiple cells. But in such a case, it is not obvious how to allocate different components to different cells. Does partitioning mitochondria into daughter cells involve any different fission operations than those that occur when a cell is not dividing? We have already defined a disconnected fission operation that acts on a cut-edge. Does there need to be a special divisional fission operation that can separate components lacking a single cut-edge?

This question of how to implement cell division in the algebraic structure raises the biological question: do mitochondria graphs in dividing cells have any discernible differences from mitochondria graphs in a non-dividing cells? Structural changes in mitochondria have been linked to changes in the cell cycle [10]. Is there any repeating motif in the mitochondria groupoid that could correspond to a complete cell cycle? Such a motif could be a natural place to impose a finite constraint on the mitochondria groupoid. Obtaining the answer to this question would require imaging a mitochondria network at different stages in the cell cycle, and then classifying what, if any, differences are observed in the corresponding mitochondrial graphs. A particularly interesting experiment is to image how a mitochondrial graph is split during cell division. Are there any definable rules that would specify or predict the cut set for the edges by which a connected component is split between the sister cells? Does division entail an imbalance between fission and fusion so as to produce more components?

#### 3.2.2 How does tip-side fusion contribute to network dynamics?

Another biological question raised by the process of constructing the mitochondria groupoid is, why does tip-side fusion exist? Since tip-to-side fusion does not affect the structure of the mitochondria groupoid, what would happen to mitochondria networks if we inhibited tip-to-side fusion? Would there be any noticeable differences? The graph from Figure 9 suggests that, in a random walk on the state-space with all transitions chosen at equal probabilities, the asymmetry created by the presence of tip-to-side fusion produces a drift towards mitochondria graphs that have a high ratio of degree-three vertices relative to degree-one vertices. We hypothesize that removing tip-to-side fusion would remove this drift. As implied by the findings of [26], this experiment could potentially be achieved by inhibiting actin-mediated processes. Alternatively, it might be possible to identify mutants affecting this process using high-content imaging combined with image analysis methods that can detect the expected changes in network morphology.

**Figure 9:**
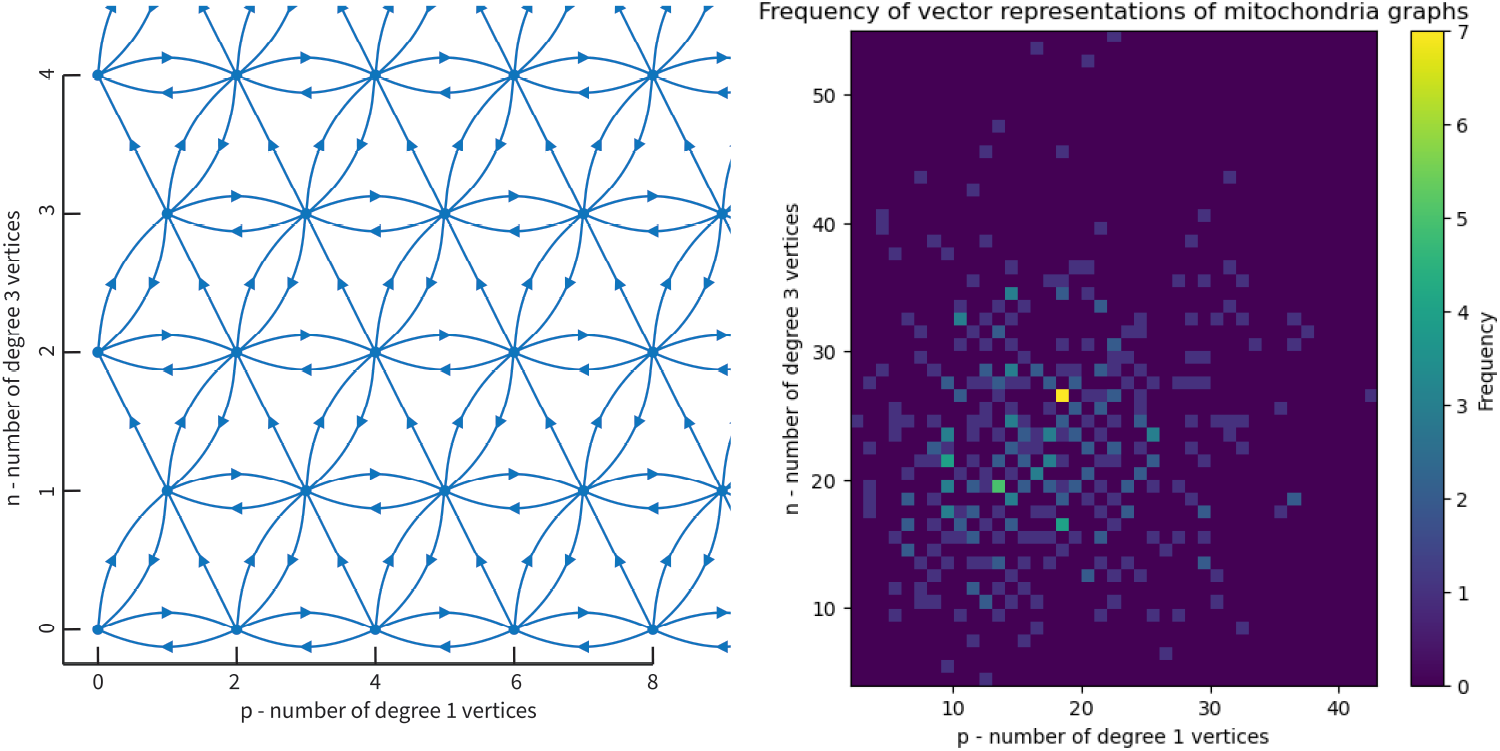
Vector groupoid *vG* forms a simplified version of the mitochondrial state-space. (Left) structure of *vG* with arrows corresponding to transitions and vertices corresponding to vectors representing collections of mitochondrial graphs with the same number of degree 1 and 3 vertices. The graph with zero vertices represents a toroidal mitochondrion. (Right) Distribution of vectors obtained from actual images of budding yeast mitochondria. This plot was generated by taking the mitograph software output provided by [18] and extracting the number of degree 1 and 3 vertices for each cell image. Frequency indicates the total number of cells for which the vector was observed in the dataset. Note that some of the vectors do not fall into the lattice due to a slight error in the mitograph software, in which branches that were near each other or overlapping in the images would be counted as vertices.

**Figure 10:**
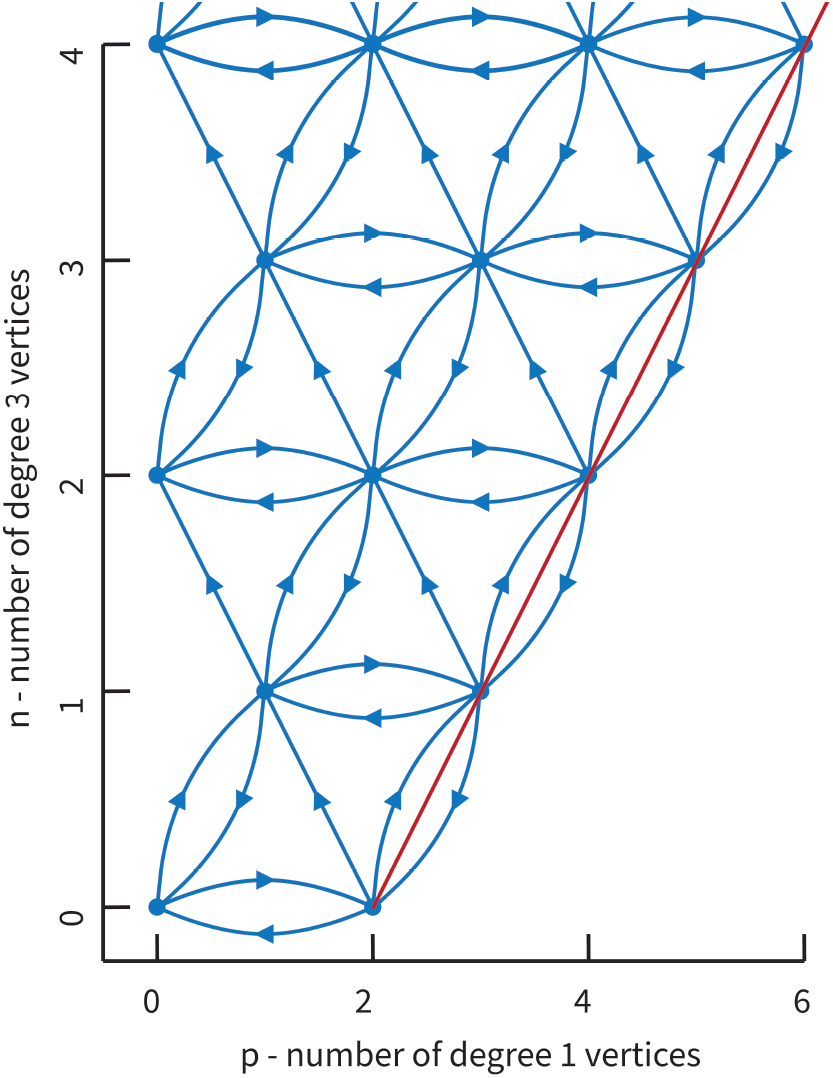
Structure of *mmG*. Dots represent the space of vertex-flip equivalence classes of single-component mitochondrial graphs, linked by transitions representing the morphological operations that generate *mmG*. Red line signifies the upper bound on p given n, corresponds to mitochondrial graphs that are trees. The left boundary at *p* = 0 corresponds to the cubic graphs, with the exception of [0, 0] being the null graph that represents a toroidal mitochondrion.

**Figure 11:**
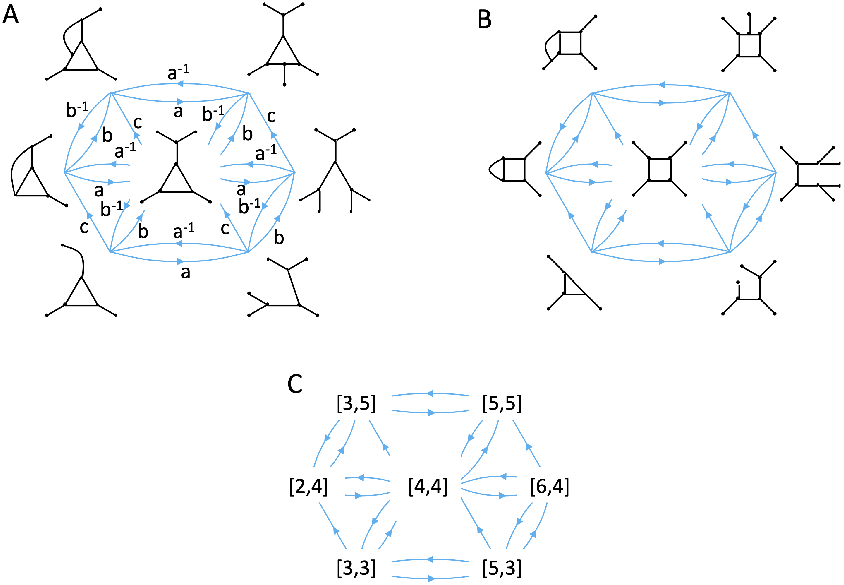
Any morphological operation applied to a pair of vertex flip-equivalent graphs yields a pair of graphs that are vertex-flip equivalent to each other.

**Figure 12:**
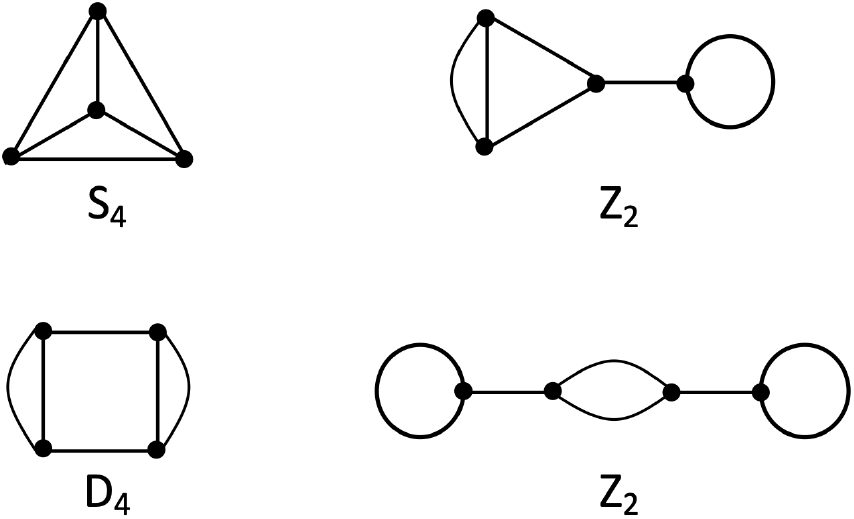
The four graphs represented by vector [4,0] with their corresponding automorphism groups.

#### 3.2.3 Why do mitochondria sometimes consist of a single giant component?

Mitochondria in budding yeast and other cell types often take the form of a single giant component together with a collection of small components, mostly I tubules ([18, 31]). As we have argued above, this empirical fact means that the single-component groupoid *mmG* can effectively represent mitochondrial dynamics in such cell types. But what biological mechanism explains this observation? Why would most of the mitochondrial material be combined together into one giant connected component, rather than dispersed among many? One could try to envision various molecular and cellular processes that might act above and beyond the mitochondrial morphological processes we have described, in order to bring about a single giant component, for example by biasing fusion to take place only in one region of the cell. However, we propose that in fact the presence of a single giant component might have a simple mathematical explanation that could explain the biological observations “for free” without the need to invoke any additional molecular mechanisms.

It has been shown that for large enough graphs with sufficient connectivity, it becomes highly likely that a single “giant component” will form. Molloy and Reed [35] establish a condition for the likely existence of a single giant component based on a weighted sum of the proportion of degrees of all vertices. Given our representation in terms of n and p, we can recast the Molloy-Reed criterion in very simple terms, such that if it is satisfied, there will be a high probability that the graph has one large component with an amount of vertices greater than or equal to a bound proportional to log(*n* + *p*). In this case, the Molloy-Reed condition is that

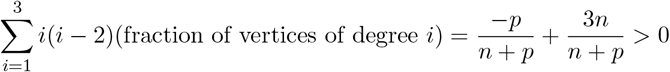

which simplifies to 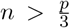. If we let *N* = *n* + *p* be the total number of vertices, and assume that each combination of *n* + *p* satisfying *n* + *p* = *N* is equally likely, then we can perform a back-of-the-envelope calculation of the probability that 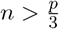

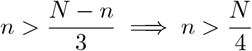

Since *n* is an integer in the range 0 ≤ *n* ≤ *N*, then the number of choices for *n* satisfying 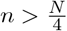 is 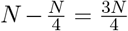. Thus, the probability that 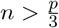 is 0.75. This assumption that each *n, p* combination is equally likely, however, is incorrect. If we instead assume that each graph with *N* vertices is equally likely, then since there is only one graph with *p* = *N* vertices, but many possible graphs with *n* ≈ *N* vertices, we conjecture that the probability that 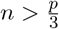 approaches 1 as *N* approaches infinity.

A key question for future research will be to ask what distribution of [*p, n*] vectors is expected from the random application of the operations with different probabilities. This is one specific aspect of a more general set of questions that we address in the next sub section.

#### 3.2.4 Is mitochondrial morphology under homeostatic control?

The morphological operations that generate the mitochondrial groupoid are physically distinct processes carried out by distinct molecular machines. Are these machines regulated by homeostatic pathways that monitor the network structure of the mitochondria? The null hypothesis is that these different processes take place at random, independently of each other or of the network structure, and each with their own rate constant. In other words the most basic assumption is that all morphological processes are strictly local in their effects and occur independently of the rest of the mitochondrial network structure. This assumption, if true, would lead to mitochondrial morphology executing a random walk in the state space defined by the mitochondrial groupoid. As these processes randomly occur, the mitochondrial morphology will randomly change, giving rise to a statistical distribution of morphologies or, in our case, a distribution of vector multisets. With the mitochondrial groupoid structure in hand, it should be possible to apply Monte Carlo methods to calculate the expected distribution of vector multisets, given the rate constants of the different processes. In many cases, the rates can be directly measured by microscopy of sufficient spatial and temporal resolution, and then plugged into the simulation. Significant deviation of the distribution observed in actual cells, from that calculated using this method, would suggest that cells may have a way to coordinate the activities of the different morphological operations so as to limit the range of morphologies that occur. This would be a form of morphological homeostasis.

If we were to find evidence for morphological homeostasis, this would open up a whole range of new biological questions that would call for mechanistic investigation. Such investigation would be impossible without a way of detecting whether the homeostasis is occurring in a given cell population, for example during a genetic screen. While one could in principle try to calculate the distribution of graphs from a set of rate constants for processes, the resulting distribution would be difficult to visualize or compare with data just because of the high complexity of the underlying space of possible graphs. The simplified representation provided by *mG, vG*, and *mmG*, provides a much more tractable framework for statistical comparisons.

On the other hand, if we were to find that the distribution of [*p, n*] vectors could in fact be explained without invoking homeostatic control, this would show that the complex structure of mitochondrial graphs may arise from a very simple set of underlying processes.

#### 3.2.5 Can we define cell states based on mitochondrial morphology?

There has been growing interest in enumerating cell types and cell states based on cluster analysis of large datasets, for example of gene expression data. The basis of such approaches is clustering - having a way to determine the distance between two cells in some kind of a state space and then clustering cells based on proximity in that space. Our algebraic representation of mitochondrial dynamics provides a way to do this, because we can quantify a “distance” based on the minimum number of operations required to convert one equivalence class to another. The approach would be to collect mitochondrial graph structures for a large number of cells, and then calculate distances between pairs of cells based on the distance calculated from the groupoid structure. One potential advantage of this approach compared to other ways to define cell state based on clustering, is that we already know a lower and upper bound on the distance, as shown in Propositions 2.5 and 2.6. Such a distance-based clustering would then lead to two general types of applications. First, the clusters would define different cell states in terms of their mitochondrial network structure. Second, the effects of genetic or metabolic perturbations could be mapped and visualized either onto the groupoid structures or onto the clusters that result, either way allowing different phenotypes to be distinguished from each other in new ways. The key to this approach is the great simplification that results in going from MG to *mG* and *mmG* based on vertex flip equivalence. Next we consider whether this approach could be generalized to other aspects of biological structure.

### 3.3 An Algebraic Approach to Cell Representation

It has long been a goal of cell biology to define cell state in some rigorous, quantifiable way, that would allow different cell types to be distinguished and related to their molecular function. One approach has been to use transcriptomics to classify cells based on gene expression patterns. However, we recognize that cell state at a purely molecular level will likely involve not only gene expression but also protein quantity, post-translational modifications, metabolites, etc. Simultaneously measuring all of these molecular scale variables in individual cells is currently a huge technological challenge, and most studies have only measured one type of variable such as transcriptomics or proteomics. Moreover, these “omics” methods are generally destructive measurements and cannot be used to measure how the system changes over time.

As an alternative strategy, it has been recognized that much of cellular function is reflected in organelle-scale cellular organization. High throughput automated microscopy, combined with increasingly powerful methods for image analysis and segmentation, have resulted in the ability to extract large amounts of quantitative data from microscopy images of cells at large scale ([36]; [37]; [38]). There has thus been growing interest in recent years in using such microscopy image data, including from live cells, to develop ways to cope with the immense complexity of molecular detail in cells, by representing cellular structure and cell state using coarser-grained representations, such as at the level of organelle morphology [3, 39, 2]. The tools currently being used to build such representations are largely drawn from statistics, using principal components analysis (PCA) and other dimensionality reduction methods to take a large number of morphological descriptors (sometimes referred to as the “morpholome”), ranging from feature descriptors like sphericity produced by standard image analysis packages to more advanced feature descriptors such as terms in a spherical harmonic expansion representing an organelle surface. By combining such measurements into a small number of modes via PCA or other dimensionality reduction methods, it is possible to produce a low dimensionality representation that can be visually comprehended and within which dynamics can be modeled in terms of a vector field [40]. Another approach that is currently seeing increased use is to employ machine learning methods such as autoencoders or GANs to learn how to encode the structure in a collection of images [41, 42, 43, 44]. This approach avoids the need to extract image features, but at the expense of interpretability compared to PCA based methods. Both of these methods have, in common, the fact that they use the images themselves, either directly or via extracted features, as the basis for defining a state space to represent cell structure.

Here we have described an alternative approach to constructing a cell representation state space. In contrast to the PCA-based morpholomic analysis based on image features or machine learning on image regions, both of which ultimately use structure as the basic element, our approach focuses on morphological processes that convert the shape of an organelle into a different shape. These operations form an algebraic structure that can be used to represent the possible states of a cell in terms of transitions between the states, and that then allows tools from abstract algebra to be brought to bear.

The challenge is that the state space, such as the space of all possible graphs, can already be very complex, and so, as with the morphological features-based methods, we need a way to reduce the complexity of the space. The approach we advocate here is to select one of the morphological processes, in our case vertex flip, and use it to construct an equivalence relation. Given this relation, we can then define a new state space in which the elements of the space are not individual shapes (for example mitochondrial graphs) but rather equivalence classes defined by the relation. This results in a vast simplification of the state space. This same approach can, in principle, be applied to any cellular structures for which an equivalence relation can be defined in terms of some set of morphological operations. By basing the reduction in space complexity on the underlying dynamics of the cellular processes, this method provides a more direct link to underlying mechanisms than would be possible with a purely statistical analysis of image features.

## 4 Summary

In conclusion, applying abstract algebra to the study of mitochondrial dynamics not only provides a formal way to represent structure as a framework for modeling or for quantifying similar states, it deepens our understanding of cellular structures and inspires experimental questions that might otherwise have remained unexplored. By conceptualizing the morphological transformation of mitochondria as elements within a groupoid, we are prompted to consider finite constraints, cyclic motifs, and the algebraic properties of mitochondrial networks in ways that traditional biological approaches might overlook. This theoretical framework thereby leads to new experimental designs that would not have emerged from a purely empirical approach.

## 5 Acknowledgements

We thank Clifford Marshall, Moumita Das, Paola Vera-Licona, Susanne Rafelski, Suliana Manley, and Anjana Badrinarayanan, for many helpful discussions. We thank Susanne Rafelski for providing the mito-dsRed construct used for yeast imaging. This work was funded by HFSP grant RGP0038/2021, NIH grant R35 GM130327, and The Center for Cellular Construction funded by NSF grant DBI1548297.

